# Predicting the subcellular location of prokaryotic proteins with DeepLocPro

**DOI:** 10.1101/2024.01.04.574157

**Authors:** Jaime Moreno, Henrik Nielsen, Ole Winther, Felix Teufel

## Abstract

Protein subcellular location prediction is a widely explored task in bioinformatics because of its importance in proteomics research. We propose DeepLocPro, an extension to the popular method DeepLoc, tailored specifically to archaeal and bacterial organisms. DeepLocPro is a multiclass subcellular location prediction tool for prokaryotic proteins, trained on experimentally verified data curated from UniProt and PSORTdb. DeepLocPro compares favorably to the PSORTb 3.0 ensemble method, surpassing its performance across multiple metrics on our benchmark experiment. The DeepLocPro prediction tool is available online at https://ku.biolib.com/deeplocpro and https://services.healthtech.dtu.dk/services/DeepLocPro-1.0/.

## Introduction

Protein subcellular location is an important aspect of protein function within cells. It is directly related to the biochemical function of proteins and it can provide valuable insights into the roles that they play in cellular processes, as well as aid in the design of biotechnological applications (McKay & Baldwin, 1990; Schiraldi et al., 2002).

While numerous machine learning (ML) methods have been developed for predicting the subcellular location of eukaryotic proteins (Almagro Armenteros et al., 2017; Blum et al., 2009; Briesemeister et al., 2009), the availability of subcellular location predictors for prokaryotic proteins is limited and not as recent (Goldberg et al., 2012; C.-S. Yu et al., 2004; N. Y. Yu et al., 2010). In recent years, Deep learning (DL) algorithms have become the method of choice for location prediction methods (Stärk et al., 2021). To this date, no DL based method for prokaryotic location prediction has been proposed.

To bridge this gap, we introduce DeepLocPro, a DL method designed specifically for predicting the subcellular location of prokaryotic proteins. DeepLocPro utilizes protein language models (pLMs), which are neural networks that have been trained on large amounts of protein sequence data to learn “the language of proteins”. By leveraging pLMs, DeepLocPro is able to capture complex patterns in proteins, such as biochemical characteristics or evolutionary information encoded in the sequences that are indicative of the final subcellular location.

DeepLocPro has been trained to work with prokaryotic proteins from a wide range of organisms covering Archaea, Gram-positive bacteria, and Gram-negative bacteria. This adaptability renders it a versatile and applicable tool across multiple fields, including microbiology and biotechnology (Drider & Rebuffat, 2011; Schiraldi et al., 2002).

## Methods

### Data

We curated experimentally verified subcellular location data from two sources: PSORTdb 4.0 (Lau et al., 2021) and UniProt release 2023_03 (The UniProt Consortium, 2023). For the PSORTdb dataset, we exclusively considered experimentally verified entries and excluded proteins with unknown IDs or those that could not be mapped to Uniprot. Our focus was on six main subcellular locations for prokaryotic proteins: cell wall and surface, extracellular space, cytoplasm, cytoplasmic membrane, outer membrane, and periplasm. We retrieved the sequences of PSORTdb entries from UniProt, resulting in a dataset of 10,241 proteins. From UniProt, we curated proteins with experimentally verified subcellular locations belonging to taxonomies Archaea (ID:2157) and Bacteria (ID:2), yielding 2,982 proteins. UniProt locations were mapped to the six main locations (Table S1). To classify bacterial data into the Gram-positive and Gram-negative categories, we followed the methodology of SignalP 6.0 (Teufel et al., 2022), which defined Gram-positive as *monoderm* phyla and Gram-negative as *diderm* (Figure 1a). Both datasets were merged, removing duplicated entries and proteins with a sequence length of fewer than 40 amino acids. Although multilabel prediction is desirable for this task (Thumuluri et al., 2022), there is not currently enough data to train the model as multilabel, as sequences with more than one location only made up a small fraction (∼3%) of the data. We therefore chose to treat the problem as a single location multi-class problem and remove the multilabel sequences from the dataset. A total of 11,970 protein sequences were obtained (Table S2).

**Figure 1.**
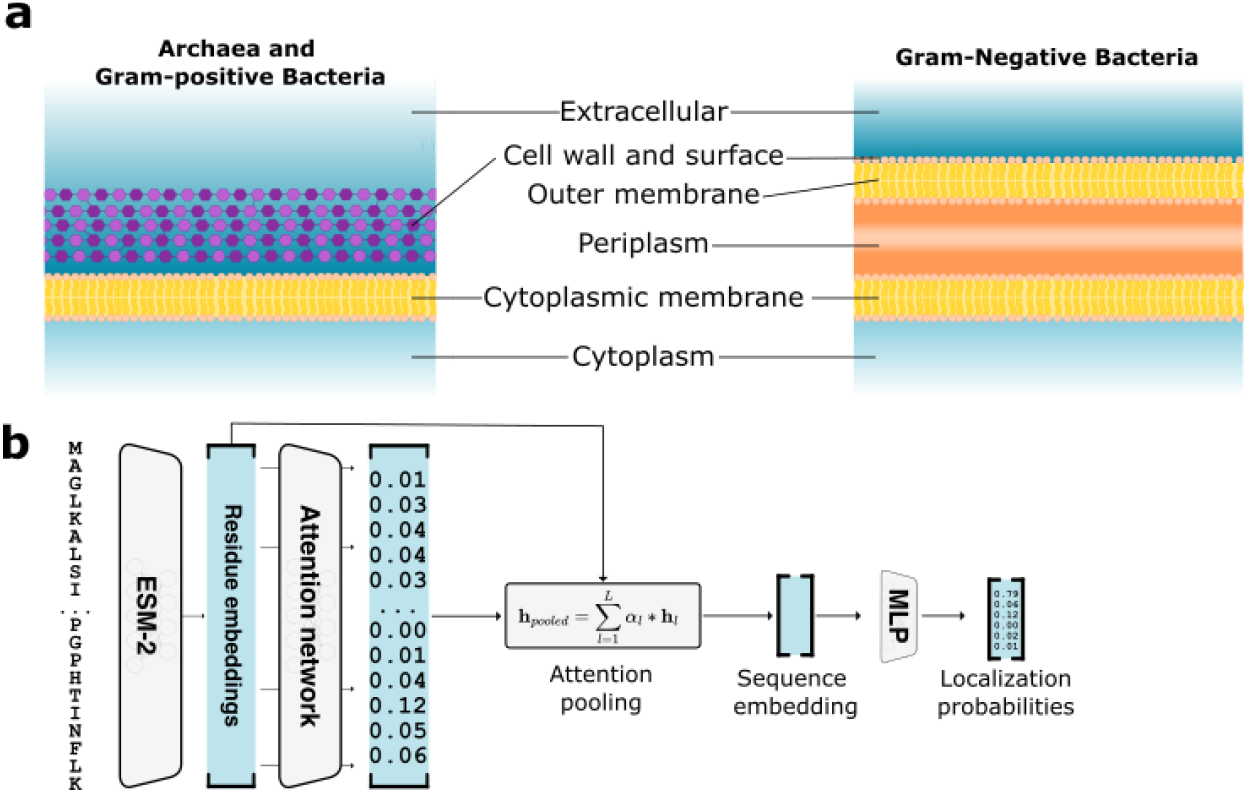
Modeling prokaryotic subcellular location. a. Prokaryotic subcellular locations for archaea, Gram-positive and Gram-negative bacteria as modeled by DeepLocPro. b. DeepLocPro model architecture. Sequences are embedded by the ESM-2 pLM, followed by an attention network that predicts a weight for each residue, which is then used to compute a weighted sum of all residue embeddings. The resulting sequence embedding serves as input to the classification layer that predicts location probabilities.

To measure model performance reliably on held out data, we homology partitioned our dataset into five folds using GraphPart (Teufel et al., 2023). Sequences in a fold have a maximum 30% Needleman-Wunsch sequence identity to any sequence in other folds. Folds were balanced for organism groups and subcellular locations using GraphPart. As a consequence, 64 proteins were removed from the dataset to achieve separation of the five folds (Table 1, Table S3).

**Table 1.** The DeepLocPro training dataset. Number of sequences per location returned by GraphPart categorized by organism group.

| Organism group | Cell wall and surface | Extracellular | Cytoplasmic | Cytoplasmic Membrane | Outer Membrane | Periplasmic | Total |
| --- | --- | --- | --- | --- | --- | --- | --- |
| Archaea | 4 | 21 | 124 | 134 | - | - | 283 |
| Negative | 21 | 581 | 4,700 | 1,793 | 756 | 566 | 8,417 |
| Positive | 62 | 475 | 2,061 | 608 | - | - | 3,206 |
| Total | 87 | 1,077 | 6,885 | 2,535 | 756 | 566 | 11,906 |

### Model

The DeepLocPro model is based on the DeepLoc 2.0 architecture (Thumuluri et al., 2022). The DeepLoc 2.0 model performs attention pooling of a protein sequence embedding, followed by a Multilayer Perceptron (MLP) for predicting class probabilities. As the embedding model, we used the publicly available pLM ESM2 with 650M parameters (Lin et al., 2023). We simplified the DeepLoc 2.0 architecture by omitting the sorting signal prediction module and training without regularization of attention weights. As DeepLocPro is a multiclass model, the sigmoidal activation function of the output layer was replaced with a Softmax (Figure 1b).

The neural network itself is agnostic to the organism group of origin, relying on the pLM’s ability to correctly infer the origin organism group from the sequence alone (Alley et al., 2019; Elnaggar et al., 2022; Teufel et al., 2022). Optionally, to avoid spurious predictions, DeepLocPro can be provided with the organism group information. If this information is provided, biologically implausible predictions (outer membrane and periplasmic proteins in archaea and gram positive organisms) are remapped to extracellular, as those locations lie beyond the inner membrane.

For training, we used Adam as the optimizer and Cross Entropy loss. We also implemented a dynamic learning rate reduction strategy, decreasing the learning rate by a factor of 0.1 when the model stopped improving for five epochs. Models were trained for a total of 60 epochs. The model was implemented in PyTorch.

### Evaluation

The model was evaluated using five-fold nested cross-validation. Having five dataset folds, each outer loop iteration involves selecting one fold as the test set, and combining the remaining four folds in different combinations for training and validation. These 4 different combinations constitute the inner loop. In total, 20 models are trained.

We optimized the model’s hyperparameters by running the model with different combinations of learning rates (0.1, 0.01, 0.001), batch sizes (8, 16, 32, 64) and dropout rates (0.1, 0.2, 0.4). For each fold, we selected the best performing hyperparameters by calculating the F1 macro average metric for the four possible inner iterations within each fold (Table S4).

After hyperparameter optimization, the final performance of the model was reported by averaging the class probabilities of each four inner loop models on their test fold. We thereby obtained a held out test prediction for each sample in our dataset. We computed performance metrics on each fold, reporting the average metric over five folds. We assessed the performance using three metrics: Accuracy, Macro F1 and Matthews Correlation Coefficient (MCC).

### Benchmark

For comparison, we report performance of PSORTb 3.0 (Yu et al., 2010), which we consider a reference method for prokaryotic subcellular location prediction due to its popularity and strong performance compared to other methods (Magnus et al., 2012). PSORTb 3.0 is an ensemble model that combines the output of multiple prediction methods. Beyond a collection of ML predictors, it also features homology-based components such as a BLAST search and sequence motifs.

While homology is a powerful mechanism for the assignment of locations to new sequences, it presents a challenge for comparing the performance of ML methods. If a test sequence is similar to the labeled data underlying the homology modules, it can be trivially classified as correct by the method. However, this performance will not be indicative of the true test performance that can be expected on new data that are dissimilar to the known training data. To mitigate such effects, we a) disabled the SCLBLAST module of PSORTb 3.0 (Supplementary Note 1) and b) created a benchmark test subset of sequences that were not available at the creation of PSORTb 3.0, using a temporal cutoff, including sequences from 2010 onwards, since it was when the model was released.

While these steps somewhat reduce performance overestimation, we note that the reported performance can still be considered overestimated, given that — contrary to the DeepLocPro evaluation — it is not possible to guarantee that the data used for testing has a maximum 30% sequence identity to the data used to train the available PSORTb 3.0 ensemble predictor. Unlike DeepLocPro, which always returns a prediction for any given protein, PSORTb 3.0 can return “Unknown” as a location prediction. For computing performance metrics, unknown predictions count as false negatives.

## Results

### DeepLocPro performance

In nested cross-validation, DeepLocPro achieved an overall accuracy of 0.92 and an overall macro F1 score of 0.81. The performance of DeepLocPro varied across different subcellular locations. We observed the highest performance on cytoplasmic and cytoplasmic membrane localized proteins, with an MCC of 0.91 and 0.88, respectively. We find that predicting locations with a small number of training samples, such as the cell wall and surface, remains challenging with an MCC of 0.59.

Stratification of performance by organism groups revealed that prediction of location in archaea is more challenging than in bacteria. The prediction of cell wall sequences shows a MCC of-0.01, indicating that DeepLocPro has random performance. We attribute this to the fact that there are only four sequences available for this category, and the archaeal cell wall being distinct from bacterial cell walls (Albers & Meyer, 2011), so that transfer learning from the more numerous bacterial sequences does not take place.

Additionally, we computed confusion matrices for all organism groups (Figure S1). We observe that distinguishing between cell wall and surface and extracellular proteins still remains challenging. This is likely due to the fact that both locations make use of the secretory pathway, while the sequence features that localize a protein to the cell wall or surface after secretion are more elusive and varied.

### Benchmarking results

On the post-2010 benchmark set, DeepLocPro outperformed PSORTb 3.0 in all three groups in terms of accuracy and macro F1 score (Table 3). Per-location performance metrics reveal that DeepLocPro outperformed PSORTb in all subcellular locations in the Gram positive and negative groups, which contains the largest number of sequences. In archaea, PSORTb surpassed DeepLocPro’s performance for extracellular proteins.

**Table 2.**
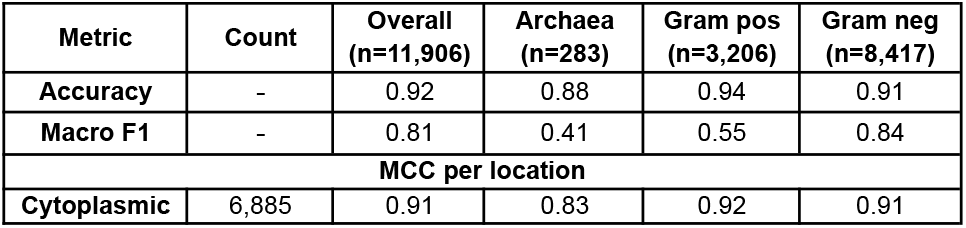
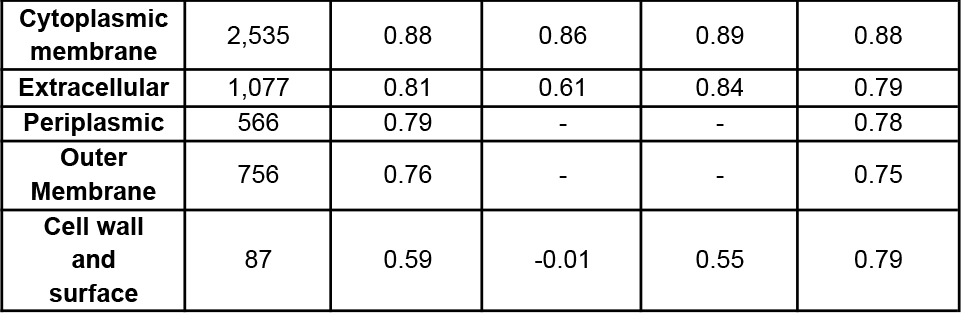
Metrics of DeepLocPro. Performance was calculated using nested cross-validation.

| Metric | Count | Overall<br>(n=11,906) | Archaea<br>(n=283) | Gram pos<br>(n=3,206) | Gram neg<br>(n=8,417) |
| --- | --- | --- | --- | --- | --- |
| **Accuracy** | - | 0.92 | 0.88 | 0.94 | 0.91 |
| **Macro F1** | - | 0.81 | 0.41 | 0.55 | 0.84 |
| **MCC per location** |  |  |  |  |  |
| **Cytoplasmic** | 6,885 | 0.91 | 0.83 | 0.92 | 0.91 |
| Cytoplasmic membrane | 2,535 | 0.88 | 0.86 | 0.89 | 0.88 |
| Extracellular | 1,077 | 0.81 | 0.61 | 0.84 | 0.79 |
| Periplasmic | 566 | 0.79 | - | - | 0.78 |
| Outer Membrane | 756 | 0.76 | - | - | 0.75 |
| Cell wall and surface | 87 | 0.59 | -0.01 | 0.55 | 0.79 |

**Table 3.** Performance of DeepLocPro and PSORTb 3.0 on the benchmark set.

| Metric | Count | Archaea (n=131) |  | Gram positive (n=366) |  | Gram negative (n=634) |  |
| --- | --- | --- | --- | --- | --- | --- | --- |
|  |  | DeepLocPro | PSORTb | DeepLocPro | PSORTb | DeepLocPro | PSORTb |
| Accuracy | - | **0.89** | 0.70 | **0.79** | 0.48 | **0.74** | 0.32 |
| Macro F1 | - | **0.37** | 0.34 | **0.44** | 0.29 | **0.75** | 0.34 |
| MCC per location |  |  |  |  |  |  |  |
| Cell wall and surface | 35 | - | - | **0.28** | -0.02 | **0.80** | 0.00 |
| Extracellular | 288 | 0.42 | **0.57** | **0.70** | 0.44 | **0.76** | 0.28 |
| Cytoplasmic | 250 | **0.81** | 0.54 | **0.67** | 0.55 | **0.68** | 0.36 |
| Cytoplasmic membrane | 325 | **0.85** | 0.65 | **0.78** | 0.51 | **0.69** | 0.59 |
| Outer membrane | 160 | - | - | - | - | **0.6** | 0.28 |
| Periplasmic | 73 | - | - | - | - | **0.72** | 0.34 |

## Discussion

The accurate prediction of subcellular location is critical for understanding the function and interactions of proteins within a cell. In this study, we developed DeepLocPro, an ML method for predicting the subcellular location of prokaryotic proteins using protein language models.

Combining experimentally verified data from two sources, we established a comprehensive dataset of over 11,900 sequences. However, for some locations such as the archaeal cell wall and surface, the number of available experimentally verified proteins is still severely limiting for training ML algorithms. While pLMs can help mitigate limited data to some extent, we find that model performance still correlates with the availability of training sequences. This indicates that future work might benefit from targeted data curation for the locations that are underrepresented in the databases. Even though comparing the performance of DeepLocPro directly to PSORTb 3.0 is challenging, as it cannot be evaluated at a sequence identity threshold of 30%, our findings indicate that DeepLocPro’s cross-validated predictions outperform PSORTb 3.0 on all but one metric, showing a strong improvement in the performance especially for bacterial proteins.

## Conclusion

We introduce DeepLocPro, a multiclass subcellular location prediction tool for prokaryotic proteins.The model is based on pre-trained protein language models that capture biochemical and evolutionary information, resulting in improved performance compared to current methods. The DeepLocPro web server is accessible online at https://ku.biolib.com/deeplocpro and https://services.healthtech.dtu.dk/services/DeepLocPro-1.0/.

