## Supplemental Data for "Predicting the subcellular location of prokaryotic proteins with DeepLocPro"

### Supplementary Data

| **UniProt location** | **Model location** | **UniProt location** | **Model location** |
| --- | --- | --- | --- |
| Cell surface | Cell wall and surface | Spore core | Cytoplasmic |
| Spore cortex | Cell wall and surface | Cellular thylakoid lumen | Cytoplasmic |
| Spore wall | Cell wall and surface | Periplasm | Periplasmic |
| Secreted | Extracellular | Forespore intermembrane space | Periplasmic |
| Host nucleus | Extracellular | Forespore inner membrane | Cytoplasmic membrane |
| Host membrane | Extracellular | Cell inner membrane | Cytoplasmic membrane |
| Host endoplasmic reticulum | Extracellular | Cellular thylakoid membrane | Cytoplasmic membrane |
| Host cytoplasm, host perinuclear region | Extracellular | Cell membrane | Cytoplasmic membrane |
| Target cell, target cell cytoplasm | Extracellular | Cell outer membrane | Outer membrane |
| Cytoplasm | Cytoplasmic | Forespore outer membrane | Outer membrane |

Table S1. **UniProt subcellular location mapping.** The original subcellular locations from Uniprot were mapped to the six location classes modeled by DeepLocPro.

| **Organism Group** | **Cell wall**  **and**  **surface** | **Extracellular** | **Cytoplasmic** | **Cytoplasmic**  **membrane** | **Outer**  **membrane** | **Periplasmic** | **Total** |
| --- | --- | --- | --- | --- | --- | --- | --- |
| **Archaea** | 4 | 21 | 124 | 138 | 0 | 0 | 287 |
| **Negative** | 21 | 588 | 4717 | 1816 | 758 | 568 | 8468 |
| **Positive** | 62 | 476 | 2064 | 613 | 0 | 0 | 3215 |
| **Total** | 87 | 1085 | 6905 | 2567 | 758 | 568 | 11970 |

Table S2. **The DeepLocPro dataset before GraphPart partitioning into five folds at 30% maximum identity.**

| **Fold** | **Cell wall**  **and**  **surface** | **Extracellular** | **Cytoplasmic** | **Cytoplasmic**  **membrane** | **Outer**  **membrane** | **Periplasmic** | **Total** |
| --- | --- | --- | --- | --- | --- | --- | --- |
| **0** | 21 | 195 | 1540 | 525 | 163 | 110 | 2554 |
| **1** | 16 | 257 | 1638 | 522 | 136 | 115 | 2684 |
| **2** | 22 | 193 | 1167 | 487 | 134 | 95 | 2098 |
| **3** | 16 | 224 | 1568 | 476 | 159 | 136 | 2579 |
| **4** | 12 | 208 | 972 | 525 | 164 | 110 | 1991 |
| **total** | 87 | 1077 | 6885 | 2535 | 756 | 566 | 11906 |

Table S3. **Compositions of the five folds of the DeepLocPro dataset.**

| **Test fold** | **Learning rate** | **Batch size** | **Drop-out** |
| --- | --- | --- | --- |
| **0** | 0.001 | 16 | 0.4 |
| **1** | 0.001 | 32 | 0.2 |
| **2** | 0.001 | 16 | 0.4 |
| **3** | 0.001 | 32 | 0.2 |
| **4** | 0.001 | 16 | 0.4 |

Table S4. **Final hyperparameters found for each nested cross-validation outer loop.**

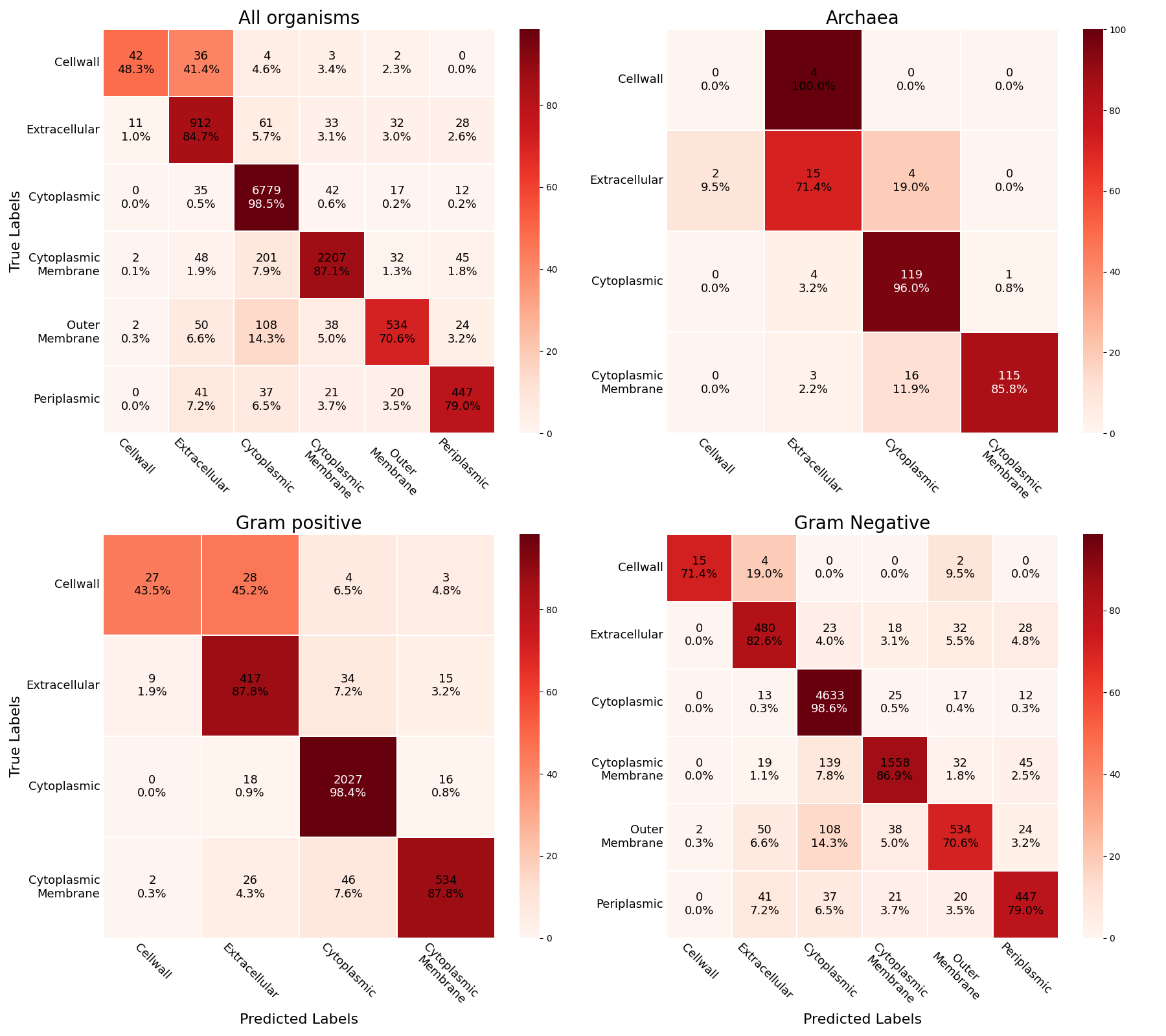
Figure S1. **Confusion matrices of DeepLocPro for each organism group**.

**Supplementary Note 1. Disabling BLAST in PSORTb 3.0**

As PSORTb 3.0 does not feature an option to disable the SCLBLAST module, we modified the brinkmanlab/psortb_commandline:1.0.2 docker image. To avoid interfering with the source code of the method, we disabled BLAST functionality by emptying the BLAST databases used by the method using the following steps.

1. Launch the docker container in interactive mode
   sudo docker run -it --entrypoint /bin/bash brinkmanlab/psortb_commandline:1.0.2
2. Navigate to /usr/local/psortb/conf/analysis/sclblast and delete the contents of the files archaea/sclblast, gramneg/sclblast and grampos/sclblast
3. Run sh makedb.sh to regenerate the databases from the sclblast files
4. Exit the docker container and use docker commit to save a modified image psortb_commandline_noblast
5. Modify the psortb perl script to execute the new image by replacing lines 11 and 61 with
   our $docker_image = "psortb_commandline_noblast";
   my $cmd = "sudo docker run --rm -v $nondocker_results_dir:$docker_results_dir -e MOUNT='$nondocker_results_dir' --entrypoint /usr/local/psortb/bin/psort -ti $docker_image /usr/local/psortb/bin/psort $extra_args -i $docker_seqfile";
